# Striosomes constrain locomotor vigor with respect to an innate valence differential

**DOI:** 10.1101/2022.05.17.492302

**Authors:** Sarah Hawes, Bo Liang, Braden Oldham, Bin Song, Lisa Chang, Da-Ting Lin, Huaibin Cai

## Abstract

Survival demands that safe and unsafe contexts are met with different locomotor profiles, yet whether striatal microcircuitry links contextual valence to spontaneous locomotor vigor is unresolved. Here we test for striosome contributions to spontaneous locomotion using ablation, calcium imaging and chemogenetics in the context of a modified Light/Dark box test. We found that locomotor speed reflects the unlearned, external valence differential present in this test, and that striosomes gate valence-oriented speed selection. Our data suggest that striosomes dampen motor vigor associated with lesser valence, or elevated anxiety.

**Highlights:**

- Mice explore the light and dark zones in Light/Dark box with different walk speed
- Striosome ablation reduces restful slowing and elevates speed relative to zone
- Many striosome neurons exhibit light-zone preference and speed related activity
- Striosome enhancement slows mice and blunts zone discrimination by speed

## Introduction

Motor control depends on striatal direct and indirect pathways’ simultaneous, complementary control over downstream nuclei (Kravitz, Freeze et al. 2010, Cui, Jun et al. 2013, Barbera, Liang et al. 2016, Yttri and Dudman 2016). Striatal projection neurons are further hypothesized to convey implicit motivation through action vigor, or speed (Dudman and Krakauer 2016), but how striatal microcircuitry achieves this is not known. A minority population of “striosome” or “patch” neurons clustered throughout the broader striatal matrix is a strong candidate for connecting external valence with implicit motivation. Striosome territories include both direct and indirect pathways similar to matrix, but striosomes’ greater connectivity to limbic structures (Gerfen 1984, Smith, Klug et al. 2016) and to dopamine neurons critical to locomotor function (Cai, Liu et al. 2014, Crittenden, Tillberg et al. 2016, Sgobio, Wu et al. 2017, Wu, Kung et al. 2019) uniquely position striosomes to integrate valence with vigor. We tested whether assessment or response to naturalistic contrasts in valence rely on striosomes using a modified Light/Dark box in combination with genetically targeted ablation, *in vivo* neuronal calcium imaging, and chemogenetics. We found that striosomes control the speed with which mice navigate the valence differential existing between high and low anxiety zones without impacting valence perception itself. This is the first evidence that striosomes regulate naturalistic motor behavior, and that in so doing, they enact implicit motivation to suit situational valence through speed selection.

## Results

### Genetic ablation of striosome neurons in Sepw1-Cre mice

To test for striosomal territories’ impact on behavior, we employed Sepw1-Cre (NP67) BAC transgenic mice in all experiments (Gerfen, Paletzki et al. 2013, Smith, Klug et al. 2016). We bilaterally injected either AAV-DIO-taCasp3 (Wu, Kung et al. 2019) to create dorsal striatal striosome ablation (SA mice), or else AAV-DIO-mCherry (controls) (**Fig. 1A**). Cre-dependent mCherry signals were colocalized with striosomal marker µ-opioid receptor (MOR1) in the dorsal striatum (**Fig. 1B**). Examination of MOR1-positive territories following behavior revealed approximately 60% loss of striosome neurons in the dorsal striatum of SA versus control mice (SA vs. control p<0.0001, **Fig. 1B-C, Supplementary Fig. S2A**). By contrast, no loss of MOR1-positive territories occurred in the ventral striatum of SA or control mice (**Fig. 1B-C**).

**Fig. 1.**
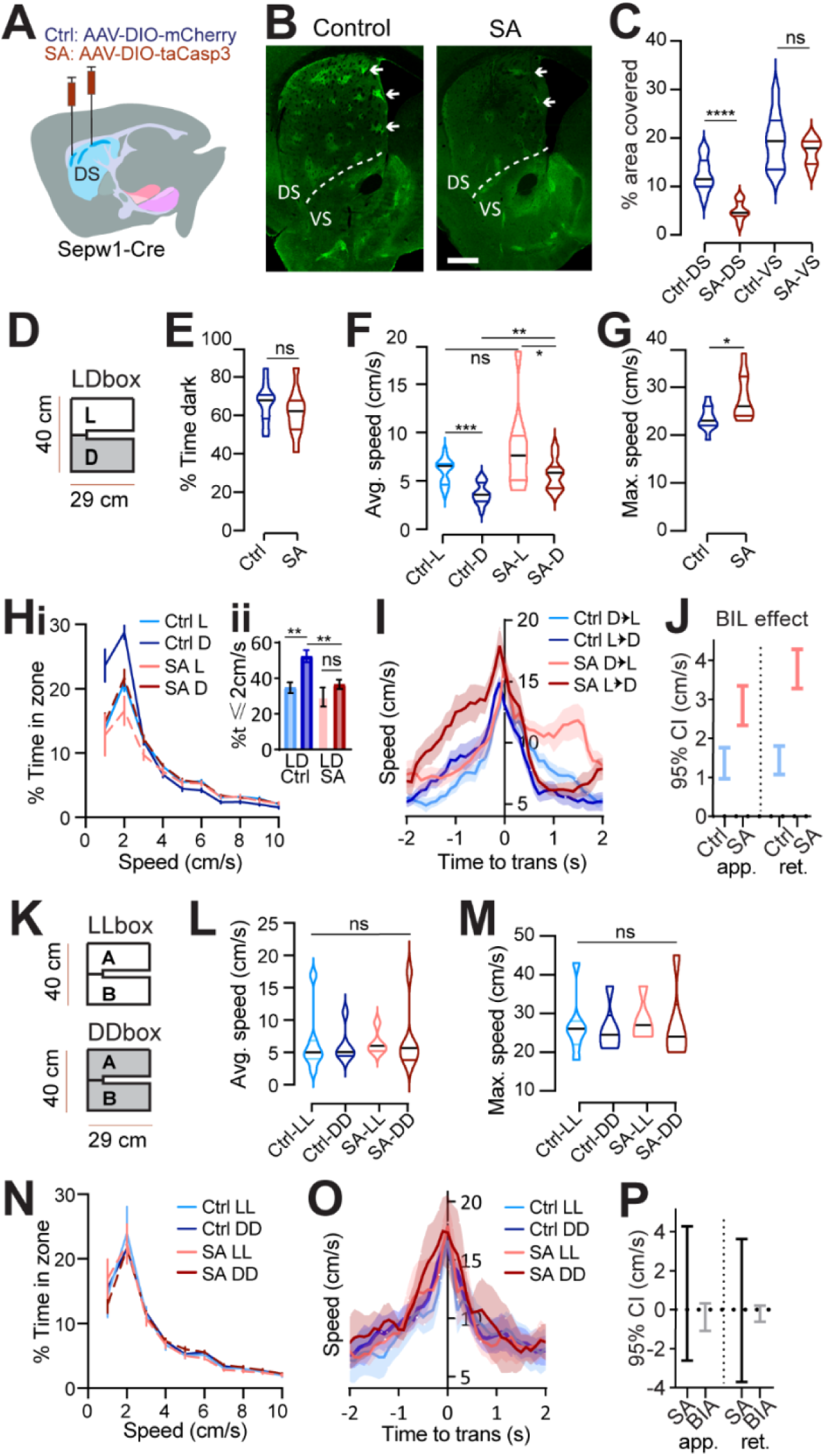
Striosome ablation reduces rest and unmasks anxious vigor at choice points. **(A)** Injection schematic. **(B)** MOR1 staining. **Scale bar: 500**μ**m. (C)** MOR1 quantification to assess ablation. 3 mice per group, n=22 Control (Ctrl) and 30 striosome ablated (SA) subregions. ^****^p<0.0001. **(D-J)** Light/Dark box. n=11 Ctrl and 10 SA mice. **(D)** LDbox schematic. **(E)** % time in dark. **(F)** Average speed. ^***^p=0.001, ^*^p=0.0371; ^**^p=0.0015 **(G)** Maximum speed. ^*^p=0.0434. **(H) i**. Speed distribution normalized to zone. **ii**. %time ≤2cm/sec L vs D Ctrl ^**^p=0.001, SA p>0.05; Ctrl vs. SA in dark ^**^p=0.0018, in light p>0.05. **(I)** Transition speed. **(J)** effect of BIL on speed is greater for SA than for Ctrls. **(K-O) LL/DD box**. n=7 Ctrl-LL, 6 Ctrl-DD, 5 SA-LL, 6 SA-DD mice. **(K)** LL/DDbox schematics. **(L)** Average speed of LL/DDbox test. **(M)** Maximum speed of LL/DDbox test. **(N)** Speed distribution normalized to side “A” of LL or DD box. For %time ≤2cm/sec, LL vs DD Ctrl p>0.05, SA p>0.05; Ctrl vs. SA in DD p>0.05, in LL p>0.05. **(O-P)** transition speed. GLME fixed effects 95% CI (cm/s) in brackets. Approach speed is unchanged by BIA [1.0937, -0.31971] or SA [-2.6119, 4.2595]. Retreat speed is unchanged by BIA [-0.62565, 0.20744] or SA [-3.708, 3.6125].

### Light/Dark box valence differential impacts speed

The Light/Dark box (LDb) test relies on the natural preference of nocturnal mice for a sheltered, dark zone despite also exploring a more anxiogenic, light zone. We differentially sub-lit a u-shaped maze to mitigate light exchange across zones (**Fig. 1D**) while also permitting uninterrupted areal video recording. Behavior in control mice verified dark preference as demonstrated by greater stay time in the dark. Like controls, SA mice demonstrated greater stay-time in the dark, indicating normal disinhibition to explore the preferred, dark zone (**Fig. 1E**). Speed was higher during exploration of the anxiogenic light zone for control and SA mice despite higher dark zone speed following ablation (L vs D control p=0.001, SA p=0.0371; control vs SA in dark p=0.0015, **Fig. 1F**). Similarity to controls indicates that striosomes are not critical for basic discrimination of situational valence.

### Striosome ablation reduces rest and unmasks anxious vigor at choice-points

While zone preference was normal, average speed in dark and maximum speed obtained were higher for SA than for control mice (p=0.0434, **Fig. 1F-G**). Importantly, the reflection of valence in speed distributions between light and dark zones present in controls was lost for SA mice, owing to a significant reduction in percent time spent at restful speeds (≤2 cm/s) in the dark among SA mice (p=0.0018, **Fig. 1H**). Given that zone discrimination and dark preference were intact, reduced stillness in the dark suggests that striosome ablation leads to failure to rest under safe conditions.

Zone transitions were accompanied by speed peaks for all mice (**Fig. 1I**), suggesting invigorated action selection at this choice-point. For controls, differential speed due to surrounding valence is immediately apparent in this moment of transition – with greater speeds expressed on either side of a transition accompanying the body-in-light (BIL, i.e., preceding a transition into dark and following a transition into light) versus body-in-dark (BID) moments (GLME fixed effects 95% CI (cm/s) in brackets. Approach speed is increased by SA [0.73218, 3.9241] and BIL [2.3911, 1.7464]. Retreat speed is increased by SA [2.2344, 2.8761] and BIL [0.89109, 3.593], **Fig. 1I**). These data support utility of the modified LDb to test an expanded LDb phenotype of valence driven locomotor performance. We next examined zone transitions among SA mice. As with controls, the shift in valence accompanying transitions was reflected at choice-points through acutely elevated speed during BIL windows for SA mice. In comparison to controls, SA transition discrimination through velocity was exaggerated thanks to selectively increased BIL vigor (GLME fixed effects 95% CI (cm/s) in brackets. SA approach [3.347, 2.3376], retreat [3.2809, 4.2788]; control approach [1.7612, 0.9698], retreat [1.0784, 1.8056], **Fig. 1I-J**). Collectively, the pattern of excess speed in SA mice suggests that striosomes naturally lower speed, supporting rest in the dark and alleviating anxious vigor at choice points.

### Speed-modulation by striosomes depends on a valence differential

LDb data implicate striosomes in lowering speed without determining if speed reduction depends fundamentally on light-level, or else on differential light-levels and the associated choice in situational valence. To distinguish these possibilities, we tested mice in chambers having the same physical dimensions as the LDb but with homogeneous illumination (**Fig. 1K**). Half the cohort was tested in a uniformly brightly lit “LLb” and the other half in a uniformly dim “DDb,” and all mice received the alternative test one-to-two-weeks later. We were surprised to find no difference distinguished LLb from DDb performance in terms of average or maximum speed for either group (**Fig. 1L-M**). Moreover, no difference between LLb and DDb speed distribution was apparent (**Fig. 1N**). Local speed peaks persisted while transitioning between equally illuminated zones, perhaps owing to the decision to turn a corner, or to pass through a relatively constricted point joining larger rooms. Yet these transition speeds did not differ between LLb and DDb (**Fig. 1O-P**, “BIA” denotes body in start side “A,” analogous to BIL as mice are started in LDb light). Thus, the side-specific LDb speed profile is not related simply to light-level but is dependent on a light versus dark valence differential. Notably, SA mice showed no elevation in speed relative to controls in LLb or DDb tests, suggesting that the impact of SA on LDb maximum speed (**Fig. 1G**) is similarly dependent on a salient valence differential. Collectively, the LLb/DDb data help to establish LDb speed as a behavioral reflection of differential situational valence and implicate striosomes in the accompanying, implicit choice in speed.

### Dark preference and greater light-zone speed are preserved during *in vivo* calcium imaging

A majority of Sepw1-Cre positive striosomes are direct-pathway neurons believed to promote locomotion (Smith, Klug et al. 2016), yet we found SA increased select, valence sensitive speed during LDb navigation. To investigate the relationship between striosome activity, valence, and speed, we analyzed *in vivo* striosomal calcium transients with genetically encoded calcium indicator GCaMP6s (Chen, Wardill et al. 2013) in 10 miniscope-mounted mice navigating the LDb (**Fig. 2A-D**). Histology following GCaMP6s transduction and behavior confirmed 1-mm diameter GRIN lens position in dorsal striatum (**Fig. 2B, Supplementary Fig. S1A-B**). Cells were identified and calcium traces (ΔF/F) were extracted (**Fig. 2C**; see **STAR Methods**) using CaImAn (Pnevmatikakis, Soudry et al. 2016, Pnevmatikakis and Giovannucci 2017, Giovannucci, Friedrich et al. 2019). Aside from a lack of separation in speeds approaching transitions (possibly due to wearing miniscope or tether), the 10 imaged mice demonstrated the LDb speed phenotype consistent with controls in our ablation experiment (**Supplementary Fig. S1C-H**).

**Fig. 2.**
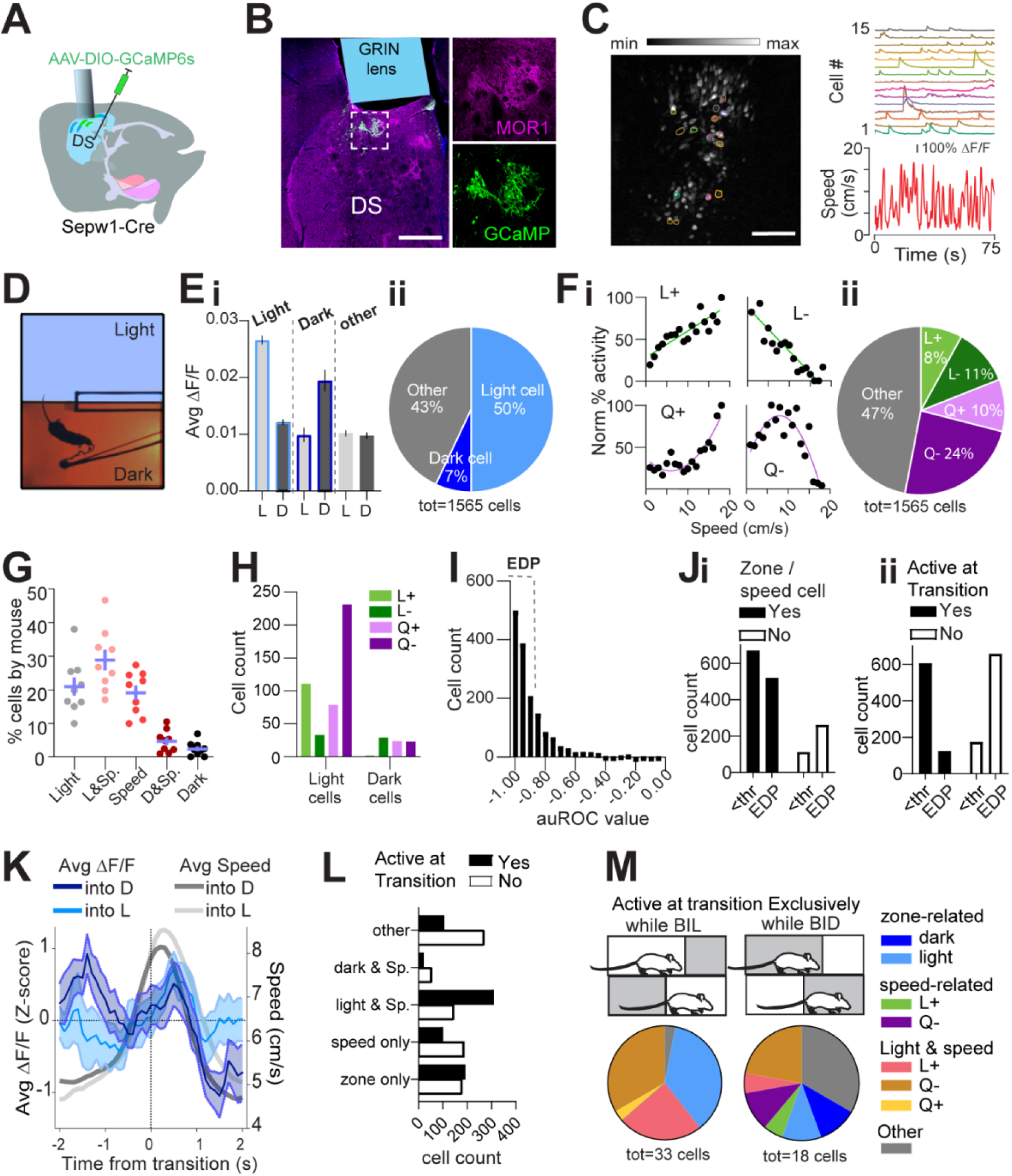
Striosome neuron activity reflects zone, speed, deceleration. Single cell calcium imaging with minniScopes. n=10 mice. **(A)** Schematic illustrating placement of GCaMP6s and GRIN lens. **(B)** Example lens placement above imaged striosomes. Selection expanded at right shows GCaMP6s and MOR1 overlap. Scale bar: 500 μm. **(C)** Left: image stack collected through GRIN lens with location of example cells circled. Scale bar: 500 μm. Right: calcium transients from example cells circled at left plotted above speed. **(D)** Light/Dark box aerial view with tethered mouse. **(E) i**. Average activity of zone-preferring (Light or Dark) and non-discriminating (Other) neurons in light “L” vs dark “D” zones. p<0.0001 Light cells; p<0.0001 Dark cells; p=0.016 Other cells. **ii**. Most zone-discriminating neurons are light-preferring. **(F) i**. Individual neurons illustrating speed relationships; clockwise from top left: linear+, linear-, quadratic+, quadratic-. **ii**. Distribution of speed relationships among imaged neurons. **(G)** Distribution of zone and/or speed related neurons is similar across all mice (n=9, excluding one mouse with fewer than 80 cells). **(H)** Speed relationships among light- and dark-preferring neurons. **(I)** Histogram of auROC. Excellent deceleration-preference “EDP” neurons are defined by |auROC| > a threshold “thr” of the auROC median value. **(J) i**. EDP neurons are under-represented among zone/speed neurons. **ii**. EDP neurons are under-represented among neurons active at zone transitions. **(K)** Overlay of average ΔF/F (blue, z-score) with mean transition speed (grey, cm/s). **(L)** Distribution of Zone/speed relationships between cells active or inactive during zone transition. **(M)** Neurons active exclusively during BIL or BID during transitions.

### Striosome neuron activity reflects zone, speed, and deceleration

We next analyzed the relationship among striosome neuron activity, zone preference, and locomotor speed. For each neuron, zone preference was determined through comparison of calcium activity in light zone to that in dark zone (see **STAR methods**). Neurons showing preferential activation in the light substantially outnumbered neurons preferentially active in the dark (**Fig. 2Ei-ii**). A recent study showed that striatal neurons display heterogeneous responses to movement speed, exhibiting both linear and nonlinear relationship (Fobbs, Bariselli et al. 2020). Similarly, we identified speed-tuned cell subtypes with both linear (positive L+ and negative L-) and quadratic nonlinear (positive Q+ and negative Q-) speed-tuning profiles (**Fig. 2Fi; STAR methods**). Approximately 53% of neurons demonstrated such speed-activity relationships (L+, L-, Q+, and Q-), with the largest fraction of speed-related cells having a quadratic negative relationship (**Fig. 2Fii**). Collectively, 76% of neurons (**Supplementary Fig. S1I**) could be defined by these relationships to zone and/or speed, and the distribution of neurons among these categories was highly consistent across mice (**Fig. 2G**). Few neurons were found to be preferentially active in dark and no preferential relationship to locomotor speed was identified among them, whereas differential speed relationships existed among light-preferring neurons (χ^2^ (3, n=532) =78.63, p<0.0001. **Fig. 2H, Supplementary Fig. S1J**). Interestingly, more than half of light-related neurons possess a Q-relationship to speed (**Fig. 2H**). These data indicate that many striosomes are engaged in speed modulation, particularly within the anxiogenic light.

We next tested for a relationship between calcium activity and change in locomotor speed. To do this we identified periods of time in which each mouse accelerated or decelerated (see STAR methods). For each neuron, we measured the activity acceleration preference using receiver operating characteristic (ROC) analysis, similar to that used in previous studies (Britten, Shadlen et al. 1992, Li, Mathis et al. 2017, Fobbs, Bariselli et al. 2020)(see STAR Methods). For a majority of striosome neurons, we found a strong relationship between deceleration and their calcium activity, represented through auROC (area under ROC curve) values close to -1 (**Fig. 2I**). We then bisected neurons into those with excellent deceleration-preference (EDP, |auROC|>median, n=783 neurons) or other neurons (|auROC|<median, n=782 neurons). Neurons with identified speed or zone relationships were less frequently EDP cells, and those for which we had been unable to identify a speed or zone relationship were disproportionately apt to be EDP cells (Fisher’s exact test, p<0.0001, **Fig. 2Ji**); this relationship persisted when dropping the threshold for EDP from the median value of 0.94 to 0.8 (**Supplementary Fig. S1Mi**). The lack of overlap sets EDP largely apart from zone or speed preferring neurons (**Supplementary Fig. S1K-L**), providing for categorization of additional neurons. As a result, we were able to ascribe a significant relationship to some combination of zone, speed, or deceleration to approximately 93% (1453 of 1565 neurons) of imaged striosomal neurons. These data suggest that a substantial subset of striosomes serves deceleration in a general sense, with no relationship to contextual valence.

### Identities of active striosomes shift with respect to zone transition

Given that striosome ablation unmasked excess locomotor vigor with respect to restful navigation in dark and during BIL transition periods, we hypothesized that striosome neurons active at these time points would possess negative correlations to speed. Limited neural activity near rest in the dark meant that we were unable to perform meaningful analysis of neuron types in these moments, and we focused instead on zone transitions. Just as EDP cells were under-represented among zone/speed related cells, they were highly under-represented among cells active at transitions (Fisher’s exact test, p<0.0001, **Fig. 2Jii**). Again, this relationship persisted despite lowing the threshold for EDP cells to 0.8 (**Supplementary Fig. S1Mii**). Thus, although we expected deceleration-related neurons to be engaged while BIL, an EDP population of striosomes appear agnostic to the decision between light and dark zones. Striosome neuron categorization by speed and zone relationships was therefore used to interrogate neural activity surrounding transitions.

A significant difference existed in striosome neuron activity surrounding transitions into light versus dark zones (GLME fixed effects 95% CI (a.u.) in brackets: ΔF/F is increased by BIL for approach [0.78166, 0.39086] and retreat [0.35765, 0.75611], **Fig. 2K, Supplementary Fig. S1O**). At face value the pattern of greater average fluorescent corresponding to periods of greater expected speed (BIL) might be taken to reflect a positive correlation between calcium transients and locomotor output. However, speed itself failed to show the expected separation upstream of zone transitions for the imaged mice (**Supplementary Fig. S1G-H**). Also, peak ΔF/F during transition toward dark occurred with BIL, but did so early in the approach, below half-maximal speeds (**Fig. 2K**). Moreover, a significant interaction between transition-active status and zone/speed relationships (χ^2^(4, n=1563) =165.2, p<0.0001, **Fig. 2L**) showed purely speed-related cells (as well as cells with no identified zone or speed relationship) were most abundant among neurons *not* active at transitions. Meanwhile, cells jointly encoding both light and speed were most abundant among transition-active neurons (**Fig. 2L, Supplementary Fig. S1N**). The most common speed relationship among transition-active neurons, including a minority exclusively active during BIL or else BID moments (**Fig. 2M**), was quadratic-negative (**Supplementary Fig. S1N**). By showing greater BIL neural activation and a shift in participating neurons toward those related to light and constrained (quadratic negative) speed, striosome neural activity at LDb zone transitions complements SA data implicating striosomes in locomotor restraint during BIL transition moments.

### Classic LDb performance is robust to acute striosome modulation by DREADDs

Given that striosome ablation enhanced valence-related speed and that calcium imaging identified striosome subsets with zone, speed, or deceleration preference, we sought to test for causality between striosomal activity and LDb speed using DREADDs for acute striosomal manipulation *in vivo*. We hypothesized that acutely suppressing striosomes would elevate anxious vigor, matching observations in SA mice, and that acutely activating striosomes would have the opposite effect – increasing restful exploration and lowering BIL speed at transitions. Using the same coordinates used for ablation, we bilaterally infused Sepw1-Cre mice with one of three viruses: AAV-DIO-hM4Di-mCherry (Gi-coupled, inhibiting), AAV-DIO-hM3Dq-mCherry (Gq-coupled, enhancing), or AAV-DIO-mCherry (control) (**Fig. 3A**). Histology confirmed colocalization of mCherry-tagged DREADDs with striosomal marker MOR1 (**Fig. 3B**). As with ablation, histological examination indicated that over half of the dorsal striosomal territories were impacted (**Fig. 3C**). Following 4-6 weeks’ recovery, we tested mice on LDb, LLb and DDb tests as before. Intraperitoneal injection of 0.3 mg/kg of DREADD-activating ligand JHU37160 (Bonaventura, Eldridge et al. 2019) preceded tests in all mice by 30-40 minutes.

**Fig. 3.**
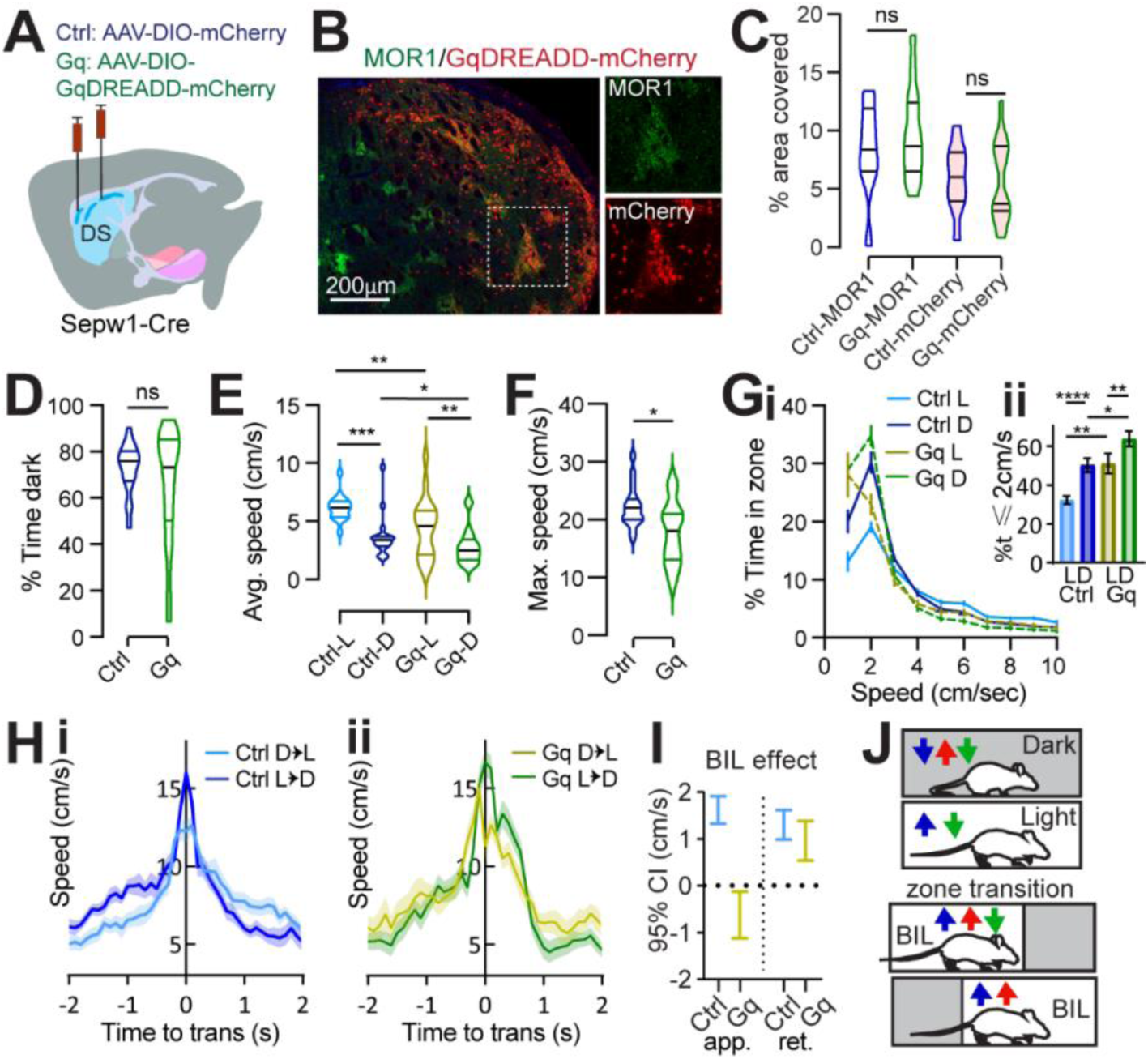
Striosome enhancement increases rest and eliminates discriminative speed at choice points. **(A)** injection schematic. **(B)** Dorsal striatal DREADD-mCherry overlaps striosomal marker MOR1. **Scale bar: 500**μμ. 20x channel separation at right. **(C)** Dorsal striatal quantification of MOR1 and mCherry in Ctrl or Gq-DREADD-transduced mice. N=one dorsal striatal subregion (from Bregma in mm: R: 0.98-1.8, M: 0.5-0.7, C: 0.02-0.26) average value from n=4-6 sections per subregion, representing 3 Ctrl and 5 Gq mice. **(D-I) Light/Dark box**. n=17 Ctrl and 20 Gq mice. **(D)** % time in dark, p>0.05. **(E)** Average speed. Ctrl-L vs Ctrl-D, ^***^p=0.0001, Gq-L vs Gq-D, p=0.0037; Ctrl-vs-Gq in Light ^**^p=0.0044, in Dark ^*^p=0.0417. **(F)** Maximum speed. ^*^p=0.01. **(G) i**. Speed distribution normalized to zone. **ii**. %time ≤2cm/sec L vs D Ctrl ^****^p<0.0001, Gq ^**^p=0.0056; Ctrl vs Gq in Light ^**^p=0.0059, in Dark ^*^p=0.0168. **(H-I)** Transition speed for controls (**Hi**) and Gq **(Hii**). **(I)** GLME by group shows speed approaching transitions is increased by BIL for controls but reduced for Gq-mice. **(J)** Cartoon summarizing speed modulation by valence under control (blue) SA (red) and Gq (green) conditions.

All groups demonstrated unaltered, basic LDb performance in that stay-time was greater in the dark (**Fig. 3D**) and speeds were higher in the light (control L vs D p=0.0001, Gq L vs D p=0.0037, **Fig. 3E**; **Supplementary Fig. S2B-C**). The average speed within zone and maximum speed obtained were lower for Gq compared to control mice (average in L p=0.0044, in p=0.0417, **Fig. 3E**, maximum p=0.01, **Fig. 3F**), demonstrating mild locomotor suppression by Gq, and unchanged for Gi mice (**Supplementary Fig. S2D)**. Overall, acute striosomal manipulation had little effect on classic measures of LDb performance. This corroborates findings from striosomal ablation, reinforcing that striosome activity does not critically determine valence perception or grossly impact affect.

### Striosome enhancement increases rest and eliminates discriminative speed at choice-points

Relative to controls, Gq-mice spent more similar time at rest (≤ 2 cm/s) in either zone, with the most pronounced increase in percent time spent at rest compared to controls occurring within the light chamber (L vs D control p<0.0001, Gq p=0.0056; control vs Gq L p=0.0059, D p=0.0168, **Fig. 3G**). Meanwhile Gi-mice showed no difference from controls in speed distribution (**Supplementary Fig. S2E**), possibly owing to already low basal activity among striosomal neurons. In terms of transition speeds, control- and Gi-mice significantly discriminated alternative zone transitions as was seen in the ablation cohort - through enhanced vigor during BIL periods (GLME fixed effects 95% CI (cm/s) in brackets: Approach speed is increased by BIL [1.0773, 0.59214]; retreat speed is increased by BIL [1.0562, 1.486], **Fig. 3Hi, Supplementary Fig. S2F**). In contrast, transition speed discrimination was impaired for Gq-mice through the apparent loss of BIL vigor (GLME fixed effects by group 95% CI (cm/s) in brackets: BIL increases approach speed for controls [1.9076, 1.3185] but decreases approach speed for Gq [-0.13168, -1.1211]. BIL increases retreat speed for controls [0.9823, 1.615] and Gq [0.53604, 1.3761], **Fig. 3Hii, 3I**). These data confirm that enhanced striosome activity restricts LDb speed. Moreover, they illustrate that striosomes suppress anxiety-related vigor, increasing time spent at restful speeds in the light, and lowering BIL speed at transitions (**Fig. 3J**).

Under homogeneous LL/DDbox conditions, no difference distinguished LLb from DDb performance for any group, and LLb and DDb performance was similar to controls for chemogenetically manipulated mice (**Supplementary Fig. S2G-N**). Whereas striosome-enhanced Gq-mice selected lower LDb speeds in the context of a valence differential, there was essentially no difference between performance of Gq-mice and controls in LLb and DDb (**Fig. 3E-G**). Therefore, as with LDb speed modulation in SA mice, LDb speed modulation by enhanced striosome activity was dependent on the presence of a valence differential.

## Discussion

By focusing on free exploration of an innate valence differential, the present work reveals a previously undescribed role for striosomes in governing implicit speed selection. A recent study parsed evaluative roles for striosome subsets and noted speed changes during conditioned place preference, but only in the absence of striosome manipulation (Xiao, Deng et al. 2020). In light of our findings, this almost certainly reflected implicit motivation governed by striosomes, analogous to the LDb control phenotype; but the impact of striosomes on speed itself remained untested. Broad striatal activity positively correlates with speed (Cui, Jun et al. 2013), yet here we find that Sepw1-defined striosomal territories, comprising approximately 85% direct and 15% indirect pathway neurons (Gerfen, Paletzki et al. 2013, Smith, Klug et al. 2016), suppress speed. This suppression of speed under naturalistic conditions is a substantial departure from often cited, correlative reports linking striosome activation due to psychomotor stimulants with hyperkinetic states (Canales and Graybiel 2000, Saka, Goodrich et al. 2004).

Our data supports and builds on recent findings that cognitive conflict and stress lead to striosomal disinhibition, and to explicit choices favoring greater risk-taking (Friedman, Homma et al. 2015, Friedman, Homma et al. 2017). We demonstrate that an innate cost-benefit choice (LDb) leads to the striosome-mediated implicit “choice” to reduce speed in riskier settings (BIL). These findings may be unified if lower anxiety-related speed manifests lower anxiety, and therefore greater risk-taking and exploration. While dark preference was robust to striosomal manipulation, future experiments are warranted to probe the important relationship between anxious locomotor vigor and anxiety itself. The lowering of BIL speed may also evince allocation of energy or attention. Striosome activity positively correlates with task-engagement and perseveration (Jenrette, Logue et al. 2019, Friedman, Hueske et al. 2020, Nadel, Pawelko et al. 2020, Nadel, Pawelko et al. 2021), implying support for habitual modes. Thus, by lowering BIL speed, striosomes may mediate a shift away from high-alert and toward habit. We believe that, by dampening response vigor to external valence, striosomes could conserve energy in safe (dark), familiar, or well-learned scenarios, alleviating cognitive demand at choice-points and under duress (Friedman, Homma et al. 2015, Beste, Muckschel et al. 2017, Beste, Muckschel et al. 2017).

The present findings may be applied to interpret or predict striosomes’ contribution to disorders of attention, mood, and motor control, including Parkinson’s disease. It will be important to continue to study striosomal modulation by diverse afferents, including the prefrontal cortex and anxiety-related basal nucleus of the stria terminalis (Smith, Klug et al. 2016, Friedman, Homma et al. 2017), and to parse striosomal impact through efferent pathways including striosome-dendron bouquets and the lateral habenula (Hong, Amemori et al. 2019, Evans, Twedell et al. 2020), in order to fully grasp striosomal importance in health and disease.

## Figure Legends

**Fig. S1.**
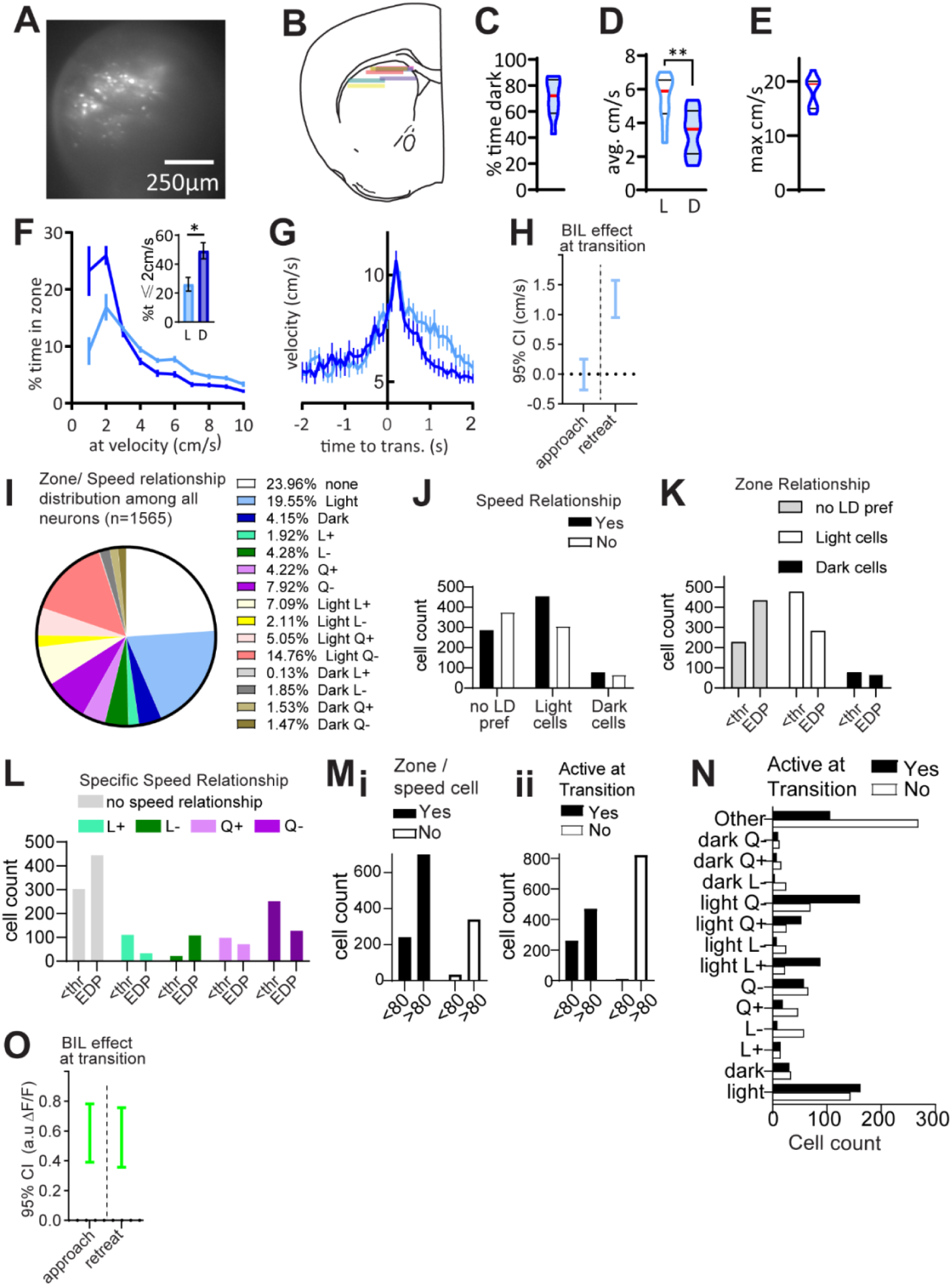
Scope-mounted Light/Dark box behavior and Ca^2+^ imaging supplemental. **(A)** example field of view through GRIN lens. **(B)** schematic of lens positions in imaged mice. **(C)** % time in dark. **(D)** average speed. Wilcoxon L vs D **p=0.0098. **(E)** maximum speed. **(F)** speed distribution normalized to zone. %time ≤2cm/sec (bar graph) Wilcoxon L vs D *p=0.0137. **(G)** transition speed. **(H)** GLME fixed effects 95% CI (cm/s) in brackets. Approach speed is unchanged by BIL [0.2506, -0.26603]. Retreat speed is increased by BIL [1.576, 0.9505]. N=10 freely moving scope-mounted mice. **(I)** distribution of zone and/or speed relationships among all recorded neurons. **(J)** A relationship exists between zone and speed encoding, X^2^(2, n=1565) =38.8, p<0.0001. **(K-L)** Excellent deceleration-prediction (EDP) cells are defined by |auROC| value above the median (threshold “thr”) of 0.94. **(K)** A relationship exists between zone- and deceleration-encoding, X^2^(2, n=1565) =114.5, p<0.0001. **(L)** A relationship exists between speed- and deceleration-encoding X^2^(2, n=1565) =172.2, p<0.0001. **(M)** Relationships between deceleration-encoding cells persist after lowering |auROC| criteria threshold to 0.8. **i**. |auROC|>0.8 neurons are disproportionately represented among zone/speed agnostic neurons. Fisher’s exact test, p<0.0001. **ii**. |auROC|>0.8 neurons are disproportionately represented among neurons not active at zone transitions. Fisher’s exact test, p<0.0001. **(N)** A relationship exists between transition-active status and zone/speed encoding. X^2^(13, n=1563) =225.1, p<0.0001. **(O)** GLME fixed effects 95% CI (a.u.) in brackets. Approach ΔF/F is increased by BIL for approach [0.78166, 0.39086] and retreat [0.35765, 0.75611].

**Fig. S2.**
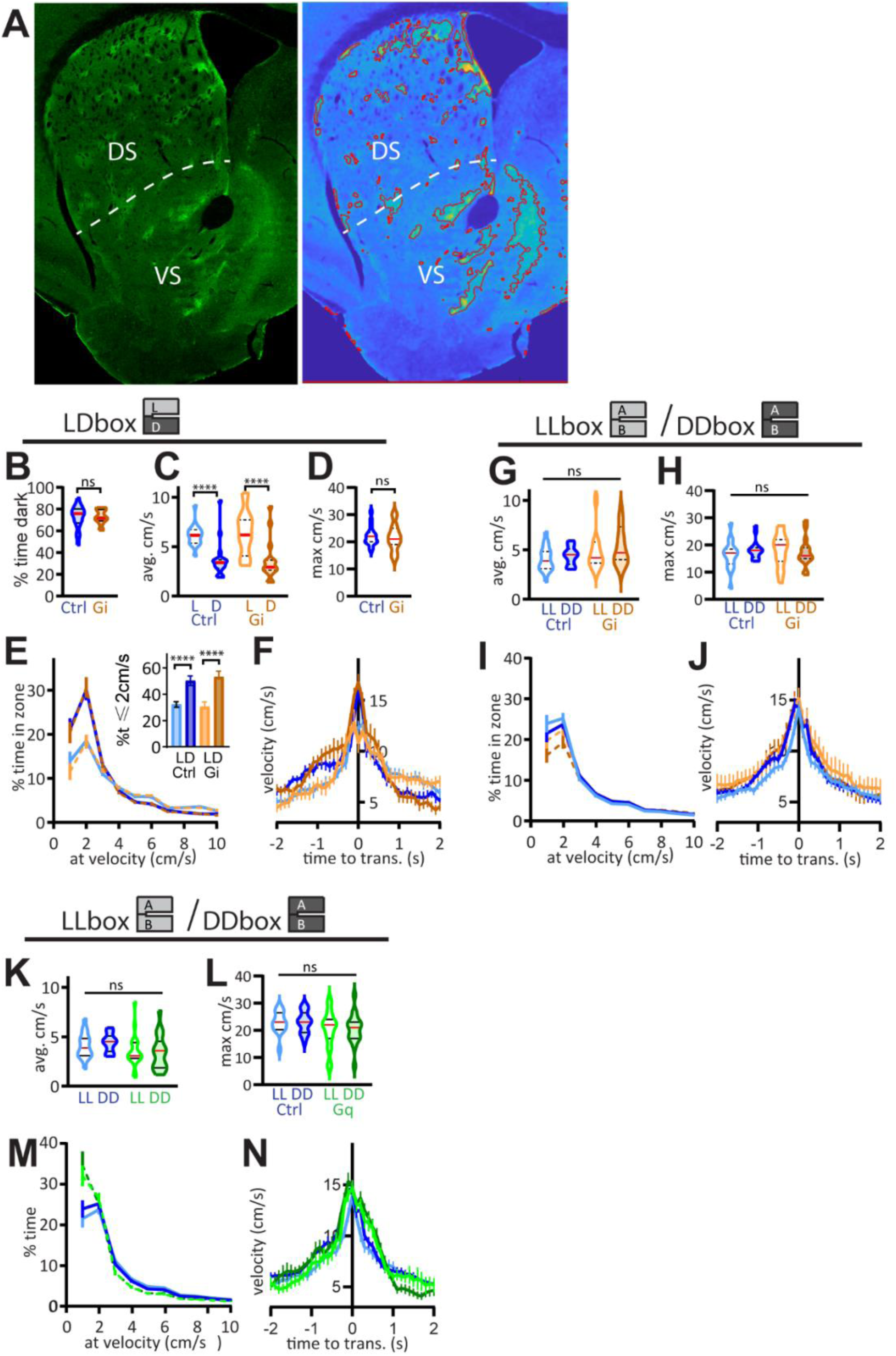
MOR1 thresholding and chemogenetics supplemental. **(A)** Striosome ablation was assessed by MOR1-positive area measurements. Example dorsal striatal (DS) and ventral striatal (VS) hemisection stained for MOR1 (left) alongside thresholded image generated in Matlab (right) used for quantifying MOR1-positive territories. **(B-F) Gi-mice L/D box**, n=17 Ctrl and 15 Gi mice. **(B)** % time in the dark. Mann-Whitney Ctrl vs Gi p=0.8232. **(C)** Average speed. 2way RM ANOVA: group F (1, 30) = 0.03006, p>0.05; light F (1, 30) = 134.3, p<0.0001. **(D)** Maximum speed. 2-tailed Mann Whitney, p=0.715. **(E)** Speed distribution normalized to zone. For %time ≤2cm/sec (bar graph) paired t-test L vs D Ctrl p<0.0001, Gi p<0.0001, unpaired t-test Ctrl vs Gi in Light p>0.05, in Dark p>0.05. **(F)** Transition speed. GLME fixed effects 95% CI (cm/s) in brackets. Approach speed is increased by BIL [1.0773, 0.59214] and unchanged by Gi [-1.0473, 1.0196]; Retreat speed is increased by BIL [1.0562, 1.486] and unchanged by Gi [-1.3062, 0.92042]. GLME by group shows approach speed is increased by BIL for control [1.9076, 1.3185] and Gi [2.2463, 1.5553]. **(G-J) Gi-mice LL/DD box**. n= mouse (16 Ctrl, 13 Gi). **(G)** Average speed. 2way RM ANOVA: lighting F (1, 27) = 3651 p=0.0667, group F (1, 27) = 2.994 p>0.05. **(H)** Maximum speed. 2way RM ANOVA: lighting F (1, 27) = 0.05506 p>0.05, group F (1, 27) = 1.402 p>0.05. **(I)** Speed distribution normalized to zone. For %time ≤2cm/sec (bar graph) Wilcoxon LL vs DD Ctrl p>0.05, Gi p=0.0105, Mann-Whitney Ctrl vs Gi in Light p>0.05, in Dark p>0.05. **(J)** Transition speed. GLME fixed effects 95% CI (cm/s) in brackets. Approach speed is slightly increased by BIA [0.85304, 0.42385], unchanged by Gi [-0.25536, 1.985]; Retreat speed is slightly increased by BIA [0.31052, 0.76904], unchanged by Gi [-0.73445, 1.7233]. **(K-N) Gq-mice LL/DD box**. n= mouse (16 Ctrl, 19 Gq). **(K)** Average speed. 2way RM ANOVA: lighting F (1, 33) = 2.676 p>0.05, group F (1, 33) = 1.343 p>0.05. **(L)** Maximum speed. 2way RM ANOVA: lighting F (1, 33) = 2.683 p>0.05, group F (1, 33) = 0.0008 p>0.05. **(M)** Speed distribution normalized to zone. For %time ≤2cm/sec Wilcoxon LL vs DD Ctrl p>0.05, Gq p>0.05, Mann-Whitney Ctrl vs Gq in LL p=0.0441, in DD p=0.0288. **(N)** Transition speed. GLME fixed effects 95% CI (cm/s) in brackets. Approach speed is increased by BIA [0.85304, 0.42385], unchanged by Gq [-1.1387, 0.89713]; Retreat speed is increased by BIA [0.31052, 0.76904], unchanged by Gq [-1.5338, 0.69961].

**Table S1.**
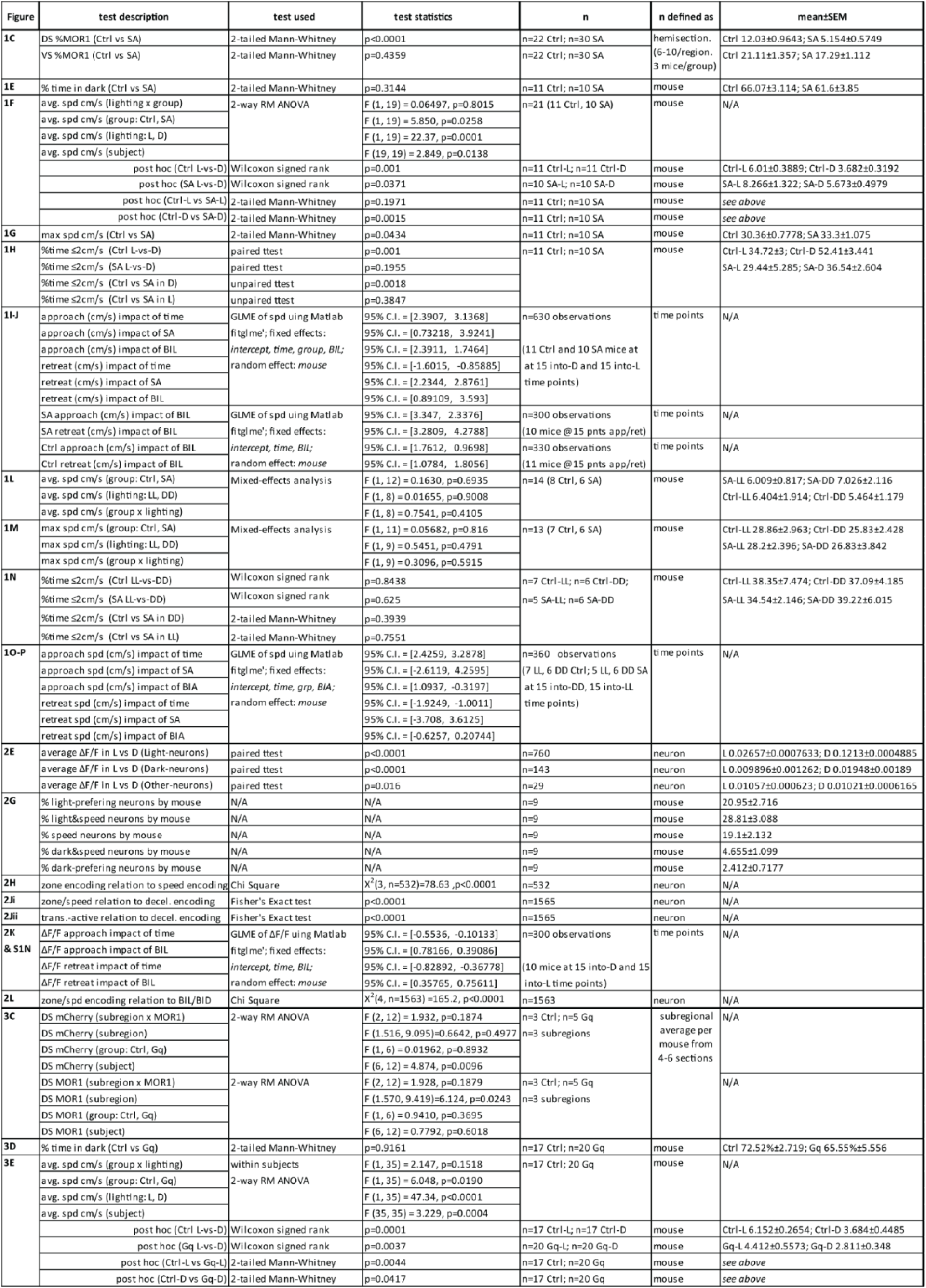
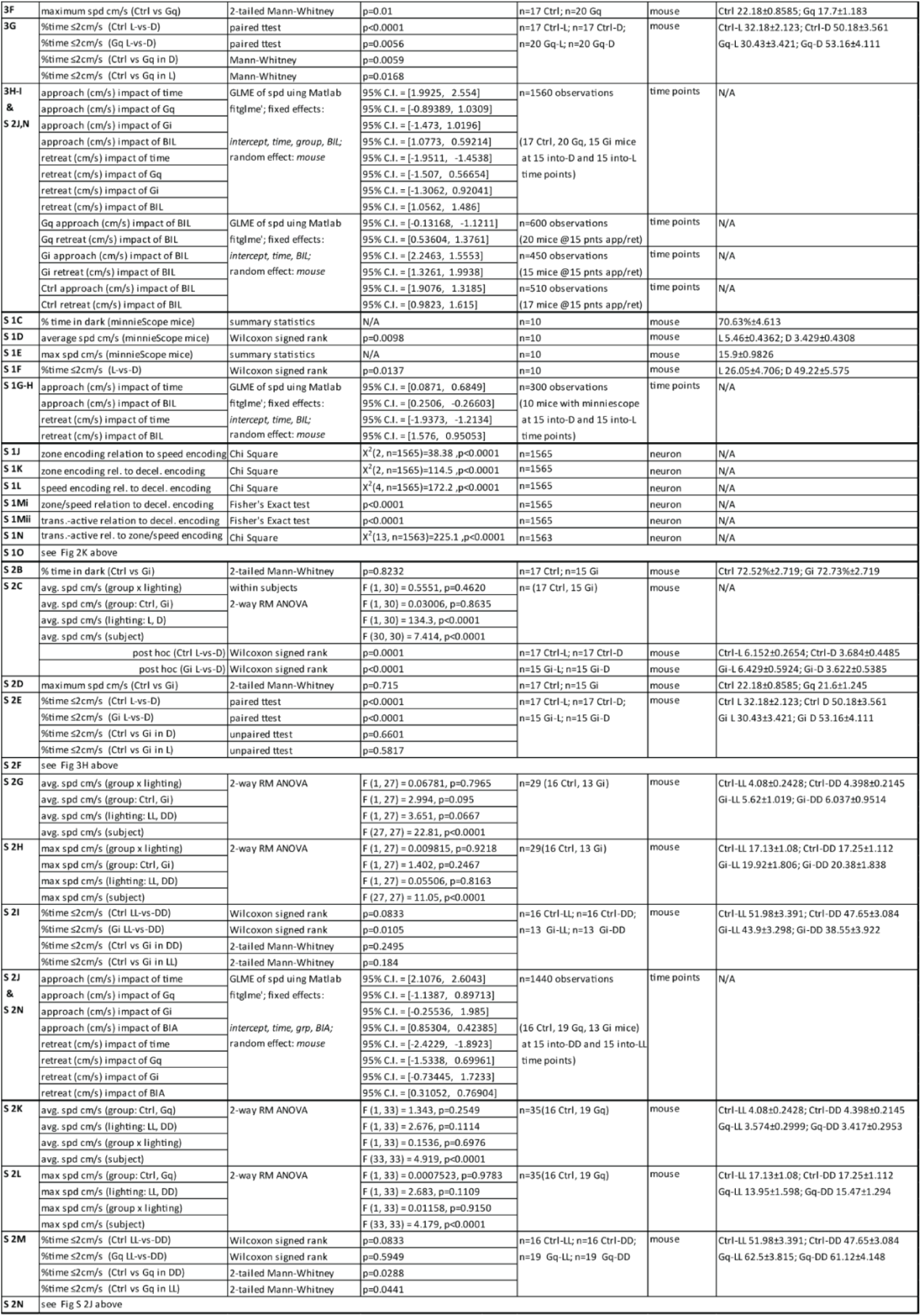

## STAR Methods

### Resource Availability

#### Key Resources Table

| REAGENT or RESOURCE | SOURCE | IDENTIFIER |
| --- | --- | --- |
| Antibodies |  |  |
| Rabbit anti-TH | Pel Freez | Cat# P40101-150, RRID: AB_461064 |
| Chicken polyclonal anti-TH | Aves Labs | Cat# TYH, RRID: AB_10013440 |
| Chicken polyclonal anti-GFP | Aves Labs | Cat# GFP-1020, RRID: AB_10000240 |
| Mouse monoclonal anti-RFP | Rockland | Cat# 200-301-379, RRID: AB_2611063 |
| Rabbit anti-MOR1 | ImmunoStar | Cat# 24216, RRID: AB_572251 |
| Alexa Fluor 488 Goat anti-Chicken IgG | Thermo Fisher | Cat# A-11039, RRID: AB_2534096 |
| Alexa Fluor 568 Donkey anti-Rabbit IgG | Thermo Fisher | Cat# A10042, RRID: AB_253401 |
| Alexa Fluor 568 Goat anti-Mouse IgG | Thermo Fisher | Cat# A-21124, RRID: AB_2535766 |
| Alexa Fluor 647 Goat anti-Rabbit IgG | Thermo Fisher | Cat# A-21244, RRID: AB_2535812 |
| Bacterial and Virus Strains |  |  |
| AAV5-EF1a-DIO-mCherry-WPRE | Vigene Biosciences | Cat. # CV17104-AV5 |
| AAV9-hSyn-DIO-hM4D(Gi)-mCherry | Addgene | Cat. # 44362-AAV9 |
| AAV9-hSyn-DIO-hM3D(Gq)-mCherry | Addgene | Cat. # 44361-AAV9 |
| AAV8-CAG-FLEX-GCaMP6s-WPRE | Vigene | Cat. # BS1-CXSAAV8 |
| AAV1-phSyn1(S)-FLEX-tdTomato-T2A-SypEGFP-WPRE | Addgene | Cat. # 51509-AAV1 |
| Chemicals, peptides, and recombinant proteins |  |  |
| Vetbond | 3M |  |
| DAPI | Thermo Fisher | Cat#: D1306 |
| Hoechst | Sigma | Cat#: 14533 |
| C&B Metabond® Quick Adhesive Cement System | Parkell | Cat#: S380 |
| JHU37160 | Hello Bio | Cat# HB6261 |
| Sodium Azide | Sigma | S2002 |
| Experimental Models:<br>Organisms/Strains |  |  |
| Mouse: SepW1-Cre [Tg(Selenow-cre)NP39Gsat/Mmucd] | Chip Gerfen, NIMH, NIH | Cat#: MMRRC_037622-UCD |
| Mouse: Drd1-flox<br>[Drd1<sup>tm2.1Stl</sup>/J] | Jackson Labs | Cat#: 025700 |
| Mouse: Drd2-flox<br>[Drd2<sup>tm1.1Mrub</sup>/J] | Jackson Labs | Cat#:020631 |
| Mouse: Lrrk2-flox | Our lab | N/A |
| Software and algorithms |  |  |
| TopScan 3.0 | CleverSys Inc. | http://cleversysinc.com; RRID:SCR_017141 |
| R | The R Project | http://www.r-project.org/; RRID:SCR_001905 |
| MATLAB | MathWorks | https://www.mathworks.com; RRID: SCR_001622 |
| Prism 8.0 | Graphpad | https://www.graphpad.com; RRID: SCR_002798 |
| ImageJ with Fiji | NIH | https://imagej.net/Fiji ; RRID: SCR_002285 |
| Miniature microscope<br>for *in vivo* calcium imaging | Barbera et al. (2016) | https://github.com/giovannibarbera/miniscope_v1.0 |
| GRIN<br>Lens AutoStereota Automated surgical instrument. | Liang et al., 2019 | https://github.com/liang-bo/AutoStereota |
| LSM 780 confocal<br>microscope | Zeiss | n/a |
| Other: |  |  |

#### Lead Contact

Additional information or requests for resources may be directed to Lead Contact Dr. Huaibin Cai.

#### Materials Availability

This study did not generate new mouse models or reagents.

#### Data and Code Availability

Raw calcium traces and MATLAB code for cell extraction are available in Git Hub https://github.com/liang-bo/StriosomeProject

## Method Details

### Mice

Animal work followed guidelines approved by Institutional Animal Care and Use Committees (IACUC) of the National Institute on Aging (NIA), NIH. Mice were male and female on a C57BL/6 background and were 3-5 months old at the time of behavioral testing, except for two females aged 6 and 8 months to meet size requirements (>25g) for undergoing GRIN lens implantation. Mice were housed in a twelve-hour light/dark cycle with ad libitum access to food and water. Behavioral tests were performed during the light cycle.

### Stereotaxic injections

Stereotaxic survival surgery was performed with aseptic technique. Anesthetized mice were head mounted to stereotaxic frames and Bregma and Lambda leveled to within 0.05 mm. Mice received infusion of AAVs (5.85×10^13 g.c./mL AAV1-FLEX-taCasp3 from Vigene, 2.42×10^13 g.c./mL AAV5-EF1a-DIO-mCherry-WPRE from Vigene, 2.1×10^13 g.c./mL AAV9-hSyn-DIO-hM4D(Gi)-mCherry from Addgene, 2.1×10^13 g.c./mL AAV9-hSyn-DIO-hM3D(Gq)-mCherry from Addgene) through a vertically held syringe (2-µL Neuros Syringe, Hamilton). Infusion of these AAVs occurred at two points bilaterally for a total of four infusion sites per mouse with the following volumes, and at the following coordinates in mm relative to Bregma: 0.7-0.8µL at AP 0.5, ML ±2.2, DV -3; 0.5-0.6µL at AP 1.5, ML ±1.8, DV -3.5. Mice receiving GCaMP6s for calcium imaging (1.3×10^13 g.c./mL AAV8-CAG-FLEX-CGaMP6s-WPRE.25.641 from Vigene) were infused into the right hemisphere only through a 1700 series Gastight Hamilton syringe with 34 gauge needle and the tip entering the head angled 30 degrees toward rostral, delivering 0.5-0.6 µL to the following coordinates relative to Bregma: AP: -0.6 mm, M/L: +2 mm, D/V: -3 mm (D/V travel at 30 degrees from vertical, to avoid scar tissue in the future tract of the GRIN implant). All infusions were controlled by a motorized stereotaxic infusion pump (Stoelting) at a rate of 50-75 nL/min, with a five-minute wait between infusion completion and needle withdrawal. Scalp was closed with sutures and a small amount of Vetbond tissue adhesive (3M), and coated in antibiotic ointment. Mice were given 5 mg/kg subcutaneous ketoprofen in lactated Ringers solution immediately following surgery, and daily for two subsequent days.

### GRIN lens implant

10-14 days following GCaMP6s infusion, mice underwent a second aseptic, stereotaxic survival surgery as detailed in (Zhang et al., 2019) to implant a 1-mm diameter Gradient-Index (GRIN) lens to dorsal striatum. Briefly, this procedure involved the precise, robot-guided, vacuum-excavation of a cylindrical tissue window centered at (relative to Bregma) A/P: +0.9mm, M/L: +2mm, and reaching -1.7 to -2mm DV from dura. In later surgeries, implant depth was guided by epifluorescence signal detected during surgery using a stereo microscope fluorescence adapter (Night Sea). After lens placement, scalp was replaced by dental cement, extending up the sides of the GRIN lens to secure the implant, and mice were administered subcutaneous ketoprofen for three days as described above. Following 4 weeks recovery, non-surgical miniscope base-mounting was performed, permitting removable placement of the miniscope prior to experiments as detailed in (Barbera et al., 2016).

### Behavior

#### Light/Dark box

The light/dark box (LDb) apparatus comprised a plexiglass chamber 40cm wide and 29cm deep, partially bisected into a U-shaped floor space with 10cm wide passage between light and dark zones (each 18 cm x 29 cm). The light zone had white walls and was sub-lit by white light; the dark had black walls and was sub-lit by red light. Translucent chamber flooring diffused light and eliminated a visual cliff. A lux meter held at the zone division and perpendicular to the floor read 500 lux toward light versus 40 lux toward dark. The test was conducted in an otherwise dark room, to which mice acclimated 30 minutes prior to testing. For testing, each mouse was gently placed into the light-zone and video recorded aerially. SA and DREADD experiments were 10 minutes. Ethanol and water were used to wipe down chamber surfaces between mice. TopScan 3.0 (CleverSys Inc.) was used for animal tracking in videos. Mice were scored as being either in the light zone or else in the dark zone, with a zone transition scored at the time (t=0) when the tail base follows the nose from one zone to the other. Speed was extracted from mouse center of mass. Maximum speed attained was defined for each mouse as the greatest speed recorded for ≥20 behavioral video frames (30 frames per second collection rate). Data exported from TopScan were organized in R and Matlab and analyzed in Prism8.

#### LL/DD box

The same chambers used for LDb were used, with either entirely red lighting and black walls (Dark/Dark box), or entirely white sub-lighting and white walls (Light/Light box). Because chambers were constructed of slick, black plastic, wall color was reversibly modified by the application or removal of adhesive dry-erase laminate. The side the mice were initially placed into was deemed Side A and opposite side (analogous to LDb dark zone in terms of order of entry) was deemed Side B.

#### Behavior with miniscope

Mice were anesthetized briefly with 2-5% isoflurane vapor in an induction chamber so that miniscopes could be head-mounted, and focus adjusted on the mouse recovering. Mice were placed in a transfer cage within the testing room and allowed to acclimate 30 minutes prior to testing. Imaging experiments occurred in six 5-minute sessions separated by 5-minute rests in the home cage with LED powered off for cooling; the light and dark sides were switched between 3^rd^ and 4^th^ sessions to control for spatial preference and to extend exploration. Otherwise, testing was completed identically to Light/Dark box described above.

### Histology and light microscopy

Anesthetized mice were transcardially perfused with ice cold phosphate buffered saline (PBS) followed by 4% paraformaldehyde (PFA, Electron Microscopy Sciences). Brains were post-fixed in 4% PFA overnight and cryoprotected in 30% Sucrose (Sigma-Aldrich). 40 µm coronal or sagittal brain slices were sectioned using Leica cryostat CM3050S (Leica Biosystems) and stored at 4°C in 0.03% Sodium Azide (Sigma-Aldrich) in PBS until use. Sections were rinsed in 0.3 M Glycine (Thermo Fisher Scientific) in PBS for 20 minutes at room temperature to quench autofluorescence, washed 3 × 5 min in PBS, and then incubated in blocking solution, rocking at 4 °C overnight. The same blocking solution was used to block and to prepare all antibody dilutions and consisted of 0.3% Triton X-100 (Sigma-Aldrich), 10% Donkey serum (Sigma-Aldrich), and 1% bovine serum albumen (Sigma-Aldrich) in PBS. Primary antibodies specific to TH (Rabbit 1:1000, Thermo Fisher Scientific, #P40101150; Chicken 1:500, Aves Labs, TYH), GFP (Chicken 1:500, Aves Labs, #GFP-1020), RFP (Mouse, Rockland Inc., #200301379) and MOR1 (Rabbit 1:3000, ImmunoStar, #24216) were freshly diluted and incubated rocking at 4°C for two nights. Following 3 × 5 min rocking in PBS, Hoechst 33342 (1:10,000, Sigma-Aldrich) or DAPI (1:10,000, Thermo Fisher Scientific, #62248) was combined in blocking solution together with secondary antibodies Alexa Fluor (Thermo Fisher Scientific) 488- (anti-chicken, #A11039), 568- (anti-Rb, #A10042; anti-Ms, #A-21124), or 647- (anti-Rb, #A21244). Secondary antibodies were diluted 1:500. Tissue was incubated in secondary antibody for one hour at room temperature, or else rocking at 4°C overnight. Fluorescence images were acquired using a laser scanning confocal microscopy LSM 780 (Zeiss). A 10x objective lens with 10% overlap in tiling was used to capture entire striatal hemispheres, while a 20x objective lens was used to capture higher magnification zones of interest.

### Calcium image processing

Calcium images were processed and analyzed using scripts in MATLAB (Mathworks). Calcium images were first stabilized using motion correction toolbox NoRMCorre (Pnevmatikakis and Giovannucci 2017). Constrained non-negative matrix factorization-based calcium image processing toolbox CaImAn-MATLAB (Pnevmatikakis, Soudry et al. 2016, Giovannucci, Friedrich et al. 2019) was used to extract calcium signals. Images of all recording sessions were concatenated and temporally sub-sampled to half for calcium signal extraction. The calcium trace (ΔF/F) was set to zero if the value was below 3× baseline noise level, and ΔF/F was then normalized by ΔF/F/max (ΔF/F) for further data analysis. For each session, a global cell map was generated, including all the extracted neurons in that session, with a neuron denoted as active in one trial if it displayed calcium transient above 3× baseline noise level. To register the spatial footprints of neurons identified across different sessions, the displacement fields of the correlation image among sessions were first calculated using MATLAB function (imregdemons) to estimate the shift due to the remounting of the miniscope and slow shift between the brain tissue and GRIN lens over time. The displacement fields were then applied to cell spatial footprints to align the rest of the sessions to the first session. Cell registration toolbox CellReg (Sheintuch, Rubin et al. 2017) was then used for cell registration. Both distance model and spatial footprint model were used. Cosine similarity was used in our data. Pair-wised cell-cell distance and spatial footprint similarity was calculated from all recording sessions of all mice to decide the threshold of whether two cells in different sessions were the same or not. P_same_ of 0.5 was used to choose the threshold. The cell with the highest spatial footprint similarity was chosen as the cell pair (same cell) if there were multiple cells that were above the threshold. Cell-to-cell mapping indices were then generated to indicate the relationship of the cell identity among sessions.

### Quantification and statistical analysis

Statistics details including description of n for all tests can be found in Supplementary Table 1. Behavior and histology data were analyzed in Prism 9.0 software (Graphpad). Data normality or non-normality was determined by D’Agostino-Pearson omnibus test before selection of parametric or non-parametric tests. Unless otherwise stated in the text, comparison between two groups was done through paired or unpaired two-tailed *t* tests, Wilcoxon signed rank or 2-tailed Mann-Whitney (as appropriate). Comparison among more than two groups was done through repeated measures ANOVA or (where missing data precluded ANOVA) through Mixed-effects analysis prior to post hoc analysis by the above-mentioned tests. Comparison of the distribution of categorical variables among neurons was done through application of Chi-square or Fisher’s exact test. Significance was determined at the p<0.05 level, and exact p-values are given for significant tests unless p<0.0001, in which case p<0.0001 is reported for brevity. Exact p-values for non-significant tests (appearing as p>0.05 for brevity in text and legends) can be found in Supplementary Table 1. Speed or ΔF/F surrounding zone transitions is analyzed -2s to -0.6s (approach) or +0.6s to +2s (retreat) from transition; significant divergence in these values is determined by non-overlap in 95% confidence intervals from generalized linear mixed-effects (GLME) models generated by Matlab function fitglme and detailed in Supplementary Table 1. Figure error bars represent standard error of the mean, and violin plots illustrate median and interquartile range.

#### Identification of light-dark zone related cells

For each cell, calcium activities in light and dark zones (ΔF/F_light_ vs ΔF/F_dark_) were compared using Mann-Whitney U test. A cell was defined as light-zone cell if ΔF/F_light_ is significantly greater than ΔF/F_dark_ (p < 0.05/n, where n is the cell number for a given mouse), and similarly a cell was defined as dark-zone cell if ΔF/F_dark_ is significantly greater than ΔF/F_light._

#### Identification of speed cells

For each cell, the Spearman correlation coefficient (SCC) between ΔF/F and locomotion speed was calculated. A cell was defined as linear+ (or linear-) cell if the SCC is significant (p < 0.05/n, where n is the cell number for a given mouse; linear+: positive SCC and linear-: negative SCC). For cells lacking a significant SCC relationship, we fitted the activity speed relationship with quadratic curve (ax^2^+bx+c). A cell was defined as quadratic+ (a>0) and quadratic- (a<0) if the goodness of fit is above 0.3.

#### Relationship between neural activity and acceleration

We first calculated the movement acceleration and binarize the acceleration by a threshold of 0.4 cm/s^2^ The acceleration-deceleration preference was measured by receiver operating characteristic (ROC) analysis (Britten, Shadlen et al. 1992, Li, Mathis et al. 2017). ROC curves were calculated by comparing the distribution of calcium activity within acceleration period versus the distribution of calcium activity within deceleration period. We then calculated the acceleration/deceleration preference by adjusted area under ROC curve (auROC) i.e., (auROC-0.5) ×2.

## Identification of transition-active cells

Those neurons active above a threshold of 0.0238 AU (determined by group activity) during 0.6 to 2 seconds preceding or 0.6 to 2 seconds following t=0 zone transition moments were defined as active during transitions.

## Acknowledgements

This work is supported in part by the Intramural Research Programs of National Institute on Aging, NIH (HC, ZIA AG000944, AG000959) and National Institute of Drug Abuse (DTL, ZIA DA000603). We thank the NIMH rodent behavioral core for assisting in behavioral tests, Dr. Charles R Gerfen of NIMH for providing the *Sepw1-Cre* mice, Shirong Lin of Fujian Provincial Hospital for help in stereotaxic surgery, and members of Cai and Lin labs for their suggestions and technical assistance.

## Author Contributions

S.H. performed stereotaxic injections and GRIN implants, conducted experiments, and analyzed data; B.L. analyzed imaging data, gave technical support; B.O. performed stereotaxic injections and histology; B.S. and L.C. performed stereotaxic injections; D.L. support in experimental design, technical support, and data analyses, S.H. and H.C. designed experiments and wrote the paper.

## Declaration of Interests

The authors declare no competing interests.

## Notes

### Competing Interest Statement

The authors have declared no competing interest.

## References

Barbera, G., B. Liang, L. Zhang, C. R. Gerfen, E. Culurciello, R. Chen, Y. Li and D. T. Lin (2016). “Spatially Compact Neural Clusters in the Dorsal Striatum Encode Locomotion Relevant Information.” Neuron 92(1): 202–213.

Beste, C., M. Muckschel, R. Rosales, A. Domingo, L. Lee, A. Ng, C. Klein and A. Munchau (2017). “Dysfunctions in striatal microstructure can enhance perceptual decision making through deficits in predictive coding.” Brain Struct Funct 222(8): 3807–3817.

Beste, C., M. Muckschel, R. Rosales, A. Domingo, L. Lee, A. Ng, C. Klein and A. Munchau (2017). “Striosomal dysfunction affects behavioral adaptation but not impulsivity-Evidence from X-linked dystonia-parkinsonism.” Mov Disord 32(4): 576–584.

Bonaventura, J., M. A. G. Eldridge, F. Hu, J. L. Gomez, M. Sanchez-Soto, A. M. Abramyan, S. Lam, M. A. Boehm, C. Ruiz, M. R. Farrell, A. Moreno, I. M. Galal Faress, N. Andersen, J. Y. Lin, R. Moaddel, P. J. Morris, L. Shi, D. R. Sibley, S. V. Mahler, S. Nabavi, M. G. Pomper, A. Bonci, A. G. Horti, B. J. Richmond and M. Michaelides (2019). “High-potency ligands for DREADD imaging and activation in rodents and monkeys.” Nat Commun 10(1): 4627.

Britten, K. H., M. N. Shadlen, W. T. Newsome and J. A. Movshon (1992). “The analysis of visual motion: a comparison of neuronal and psychophysical performance.” J Neurosci 12(12): 4745–4765.

Cai, H., G. Liu, L. Sun and J. Ding (2014). “Aldehyde Dehydrogenase 1 making molecular inroads into the differential vulnerability of nigrostriatal dopaminergic neuron subtypes in Parkinson’s disease.” Transl Neurodegener 3: 27.

Canales, J. J. and A. M. Graybiel (2000). “A measure of striatal function predicts motor stereotypy.” Nat Neurosci 3(4): 377–383.

Chen, T. W., T. J. Wardill, Y. Sun, S. R. Pulver, S. L. Renninger, A. Baohan, E. R. Schreiter, R. A. Kerr, M. B. Orger, V. Jayaraman, L. L. Looger, K. Svoboda and D. S. Kim (2013). “Ultrasensitive fluorescent proteins for imaging neuronal activity.” Nature 499(7458): 295–300.

Crittenden, J. R., P. W. Tillberg, M. H. Riad, Y. Shima, C. R. Gerfen, J. Curry, D. E. Housman, S. B. Nelson, E. S. Boyden and A. M. Graybiel (2016). “Striosome-dendron bouquets highlight a unique striatonigral circuit targeting dopamine-containing neurons.” Proc Natl Acad Sci U S A 113(40): 11318–11323.

Cui, G., S. B. Jun, X. Jin, M. D. Pham, S. S. Vogel, D. M. Lovinger and R. M. Costa (2013). “Concurrent activation of striatal direct and indirect pathways during action initiation.” Nature 494(7436): 238–242.

Dudman, J. T. and J. W. Krakauer (2016). “The basal ganglia: from motor commands to the control of vigor.” Curr Opin Neurobiol 37: 158–166.

Evans, R. C., E. L. Twedell, M. Zhu, J. Ascencio, R. Zhang and Z. M. Khaliq (2020). “Functional Dissection of Basal Ganglia Inhibitory Inputs onto Substantia Nigra Dopaminergic Neurons.” Cell Rep 32(11): 108156.

Fobbs, W. C., S. Bariselli, J. A. Licholai, N. L. Miyazaki, B. A. Matikainen-Ankney, M. C. Creed and A. V. Kravitz (2020). “Continuous Representations of Speed by Striatal Medium Spiny Neurons.” J Neurosci 40(8): 1679–1688.

Friedman, A., D. Homma, B. Bloem, L. G. Gibb, K. I. Amemori, D. Hu, S. Delcasso, T. F. Truong, J. Yang, A. S. Hood, K. A. Mikofalvy, D. W. Beck, N. Nguyen, E. D. Nelson, S. E. Toro Arana, R. H. Vorder Bruegge, K. A. Goosens and A. M. Graybiel (2017). “Chronic Stress Alters Striosome-Circuit Dynamics, Leading to Aberrant Decision-Making.” Cell 171(5): 1191–1205 e1128.

Friedman, A., D. Homma, L. G. Gibb, K. Amemori, S. J. Rubin, A. S. Hood, M. H. Riad and A. M. Graybiel (2015). “A Corticostriatal Path Targeting Striosomes Controls Decision-Making under Conflict.” Cell 161(6): 1320–1333.

Friedman, A., E. Hueske, S. M. Drammis, S. E. Toro Arana, E. D. Nelson, C. W. Carter, S. Delcasso, R. X. Rodriguez, H. Lutwak, K. S. DiMarco, Q. Zhang, L. I. Rakocevic, D. Hu, J. K. Xiong, J. Zhao, L. G. Gibb, T. Yoshida, C. A. Siciliano, T. J. Diefenbach, C. Ramakrishnan, K. Deisseroth and A. M. Graybiel (2020). “Striosomes Mediate Value-Based Learning Vulnerable in Age and a Huntington’s Disease Model.” Cell 183(4): 918-934.e949.

Gerfen, C. R. (1984). “The neostriatal mosaic: compartmentalization of corticostriatal input and striatonigral output systems.” Nature 311(5985): 461–464.

Gerfen, C. R., R. Paletzki and N. Heintz (2013). “GENSAT BAC cre-recombinase driver lines to study the functional organization of cerebral cortical and basal ganglia circuits.” Neuron 80(6): 1368–1383.

Giovannucci, A., J. Friedrich, P. Gunn, J. Kalfon, B. L. Brown, S. A. Koay, J. Taxidis, F. Najafi, J. L. Gauthier, P. Zhou, B. S. Khakh, D. W. Tank, D. B. Chklovskii and E. A. Pnevmatikakis (2019). “CaImAn an open source tool for scalable calcium imaging data analysis.” Elife

Hong, S., S. Amemori, E. Chung, D. J. Gibson, K. I. Amemori and A. M. Graybiel (2019). “Predominant Striatal Input to the Lateral Habenula in Macaques Comes from Striosomes.” Curr Biol 29(1): 51–61 e55.

Jenrette, T. A., J. B. Logue and K. A. Horner (2019). “Lesions of the Patch Compartment of Dorsolateral Striatum Disrupt Stimulus-Response Learning.” Neuroscience 415: 161–172.

Kravitz, A. V., B. S. Freeze, P. R. Parker, K. Kay, M. T. Thwin, K. Deisseroth and A. C. Kreitzer (2010). “Regulation of parkinsonian motor behaviours by optogenetic control of basal ganglia circuitry.” Nature 466(7306): 622–626.

Li, Y., A. Mathis, B. F. Grewe, J. A. Osterhout, B. Ahanonu, M. J. Schnitzer, V. N. Murthy and C. Dulac (2017). “Neuronal Representation of Social Information in the Medial Amygdala of Awake Behaving Mice.” Cell 171(5): 1176–1190 e1117.

Nadel, J. A., S. S. Pawelko, D. Copes-Finke, M. Neidhart and C. D. Howard (2020). “Lesion of striatal patches disrupts habitual behaviors and increases behavioral variability.” PLoS One 15(1): e0224715.

Nadel, J. A., S. S. Pawelko, J. R. Scott, R. McLaughlin, M. Fox, M. Ghanem, R. van der Merwe, N. G. Hollon, E. S. Ramsson and C. D. Howard (2021). “Optogenetic stimulation of striatal patches modifies habit formation and inhibits dopamine release.” Sci Rep 11(1): 19847.

Pnevmatikakis, E. A. and A. Giovannucci (2017). “NoRMCorre: An online algorithm for piecewise rigid motion correction of calcium imaging data.” J Neurosci Methods 291: 83–94.

Pnevmatikakis, E. A., D. Soudry, Y. Gao, T. A. Machado, J. Merel, D. Pfau, T. Reardon, Y. Mu, C. Lacefield, W. Yang, M. Ahrens, R. Bruno, T. M. Jessell, D. S. Peterka, R. Yuste and L. Paninski (2016). “Simultaneous Denoising, Deconvolution, and Demixing of Calcium Imaging Data.” Neuron 89(2): 285–299.

Saka, E., C. Goodrich, P. Harlan, B. K. Madras and A. M. Graybiel (2004). “Repetitive behaviors in monkeys are linked to specific striatal activation patterns.” J Neurosci 24(34): 7557–7565.

Sgobio, C., J. Wu, W. Zheng, X. Chen, J. Pan, A. G. Salinas, M. I. Davis, D. M. Lovinger and H. Cai (2017). “Aldehyde dehydrogenase 1-positive nigrostriatal dopaminergic fibers exhibit distinct projection pattern and dopamine release dynamics at mouse dorsal striatum.” Sci Rep 7(1): 5283.

Sheintuch, L., A. Rubin, N. Brande-Eilat, N. Geva, N. Sadeh, O. Pinchasof and Y. Ziv (2017). “Tracking the Same Neurons across Multiple Days in Ca(2+) Imaging Data.” Cell Rep 21(4): 1102–1115.

Smith, J. B., J. R. Klug, D. L. Ross, C. D. Howard, N. G. Hollon, V. I. Ko, H. Hoffman, E. M. Callaway, C. R. Gerfen and X. Jin (2016). “Genetic-Based Dissection Unveils the Inputs and Outputs of Striatal Patch and Matrix Compartments.” Neuron 91(5): 1069–1084.

Wu, J., J. Kung, J. Dong, L. Chang, C. Xie, A. Habib, S. Hawes, N. Yang, V. Chen, Z. Liu, R. Evans, B. Liang, L. Sun, J. Ding, J. Yu, S. Saez-Atienzar, B. Tang, Z. Khaliq, D. T. Lin, W. Le and H. Cai (2019). “Distinct Connectivity and Functionality of Aldehyde Dehydrogenase 1a1-Positive Nigrostriatal Dopaminergic Neurons in Motor Learning.” Cell Rep 28(5): 1167–1181 e1167.

Xiao, X., H. Deng, A. Furlan, T. Yang, X. Zhang, G. R. Hwang, J. Tucciarone, P. Wu, M. He, R. Palaniswamy, C. Ramakrishnan, K. Ritola, A. Hantman, K. Deisseroth, P. Osten, Z. J. Huang and B. Li (2020). “A Genetically Defined Compartmentalized Striatal Direct Pathway for Negative Reinforcement.” Cell 183(1): 211–227 e220.

Yttri, E. A. and J. T. Dudman (2016). “Opponent and bidirectional control of movement velocity in the basal ganglia.” Nature 533(7603): 402–406.

